# Inter- and Intra-Generation Genomic Predictions for Douglas-fir Growth in Unobserved Environments

**DOI:** 10.1101/540765

**Authors:** Blaise Ratcliffe, Francis Thistlethwaite, Omnia Gamal El-Dien, Eduardo P. Cappa, Ilga Porth, Jaroslav Klápště, Charles Chen, Tongli Wang, Michael Stoehr, Yousry A. El-Kassaby

## Abstract

Conifers are prime candidates for genomic selection (GS) due to their long breeding cycles. Previous studies have shown much reduced prediction accuracies (PA) of breeding values in unobserved environments, which may impede its adoption. The impact of explicit environmental heterogeneity modeling including genotype-by-environment (G×E) interaction effects using environmental covariates (EC) in a reaction-norm genomic prediction model was tested using single-step GBLUP (ssGBLUP). A three-generation coastal Douglas-fir experimental population with 14 genetic trials (*n* = 13,615) permitted estimation of intra- and inter-generation PA in unobserved environments using 66,969 SNPs derived from exome capture. Intra- and inter-generation PAs ranged from 0.447-0.640 and 0.317-0.538, respectively. The inclusion of ECs in the prediction models explained up to 23% of the phenotypic variation for the fully specified model and resulted in the best model fit. Modeling G×E effects in the training population increased PA up to 6% and 13% over the base model for inter- and intra-generations, respectively. GS-PA can be substantially improved using ECs to explain environmental heterogeneity and G×E effects. The ssGBLUP methodology allows historical genetic trials containing non-genotyped samples to contribute in genomic prediction, and, thus, effectively boosting training population size which is a critical step. Further pheno- and enviro-typing developments may improve GS-PA.

## INTRODUCTION

Breeding conifer species for phenotypic improvement is challenged due to late expression of important phenotypes related to productivity and their late sexual maturity, causing extensive recurrent selection cycles. Genomic selection (GS) can address such shortcomings through early prediction of phenotypes based on large numbers of jointly considered genomic markers, most commonly, single nucleotide polymorphisms (SNPs) (Meuwissen *et al*. 2001). This solution is realized through improved management of co-ancestry and increased genetic gain via enhanced precision and accuracy of pairwise kinship estimates for the breeding population. The recent exploration of GS within the forest tree breeding framework has produced numerous promising studies, indicating that GS prediction accuracies are able to at least match and often surpass pedigree-based predictions (see reviews by Grattapaglia 2017; Grattapaglia *et al*. 2018).

In forest trees, the effect of genotype-by-environment (G×E) interactions can be considerable and, if overlooked, detrimental to genetic gain optimization. Therefore, it is commonplace to regionalize tree breeding efforts based on available biogeoclimatic information to avoid maladaptation of deployed stock (*i.e*., defining species’ breeding zones) (Burdon 1977). However, maladaptation may still occur within regionalized areas due to trees’ long lifespans, their placement within highly heterogeneous growing environments, along with ongoing and predicted shifting of climatic boundaries due to climate change. Thus, it is essential to account for the effect of G×E interactions when predicting breeding values to either select genotypes that are stable across the breeding region or match specific genotypes with particular locations to capture additional gain (Li *et al*. 2017) in the future. In forest trees’ genetic evaluations, G×E interactions are typically modeled using mixed linear models that allow for covariances to be estimated between environments assuming that a character measured in two environments represents two distinct traits (*i.e*., Type-B genetic correlations; Burdon 1977) (e.g. see e Silva *et al*. 2005; Cappa *et al*. 2012; Bian *et al*. 2014). However, for large numbers of environments, typically, this approach is computationally not practical and simplifications of covariance structures or factor analytic models must be put in place (Isik *et al*. 2017).

Recent GS studies in conifers have demonstrated that predictions *across* environments suffer from marked decreases in accuracy when the prediction model’s training population does not include phenotypic observations from the environment to which genomic predictions were targeted (Gamal El-Dien *et al*. 2015; Thistlethwaite *et al*. 2017; Chen *et al*. 2018). Consequently, estimated marker effects would be considered specific to the environment(s) of the training populations. Without phenotypic observations from the target environment, genomic prediction is challenging. In crop plant systems, high-dimensional environmental covariates (ECs) have been implemented in GS models to improve multi-environment genomic predictions (Jarquín *et al*. 2014). The reaction-norm models use covariance structures analogous to the realized relationship matrix (VanRaden 2008) to model environmental and G×E interaction effects. Pérez-Rodríguez *et al*. (2015) and Morais Júnior *et al*. (2018) extended the use of the reaction-norm models to include the average numerator relationship matrix (***A***) and the single-step combined relationship matrix (***H***), respectively. Pedigree-based reaction-norm methods such as random regression were only very recently proposed in a forest tree G×E interactions study (Marchal *et al*. 2019).

Single-step genomic evaluation (ssGBLUP) is a unified approach that allows the incorporation of phenotypic, genomic, and pedigree information into a single analysis (Legarra *et al*. 2009; Misztal *et al*. 2009). This methodology allows the prediction of breeding values for genotyped and non-genotyped individuals to be on the same scale, avoiding bias and complex multi-step analyses (Vitezica *et al*. 2011). It also allows for the phenotypes of non-genotyped individuals to participate in the estimation of marker effects, effectively boosting the accuracy of prediction. Thus, the method also provides a cost-effective entry into GS as relatively few important individuals can be genotyped while phenotypic records of rogued trials can also easily be implemented yielding a modern analysis for estimating marker effects or breeding values. The use of ssGBLUP has recently been demonstrated to be effective in genetic evaluations of *Picea glauca* (Ratcliffe *et al*. 2017), *Eucalyptus grandis* (Cappa *et al*. 2018), and *Eucalyptus nitens* (Klápště *et al*. 2018). However, ssGBLUP-based reaction-norm remains untested in outcrossing species such as forest trees.

A limitation of the ssGBLUP method, however, is that the combined genetic relationship matrix (***H***) can be very dense when evaluating many individuals, leading to lengthy computation times (Legarra and Ducrocq 2012). Wang *et al*. (2012a) demonstrated that genomic estimated breeding values (GEBV) of the genotyped individuals from ssGBLUP analyses can be used to calculate SNP marker effects. Lourenco *et al*. (2015) applied this approach to American Angus cattle, coining it ‘indirect prediction’. The indirect prediction approach improves computational efficiency and allows for fast prediction of GEBV for new genotyped trees via SNP marker effects as opposed to a full ssGBLUP evaluation where new genotyped individuals need to be explicitly included.

This study is based on the ‘maritime low’ coastal Douglas-fir (CDF) (*Pseudotsuga menziesii* (*Mirb*.) Franco var. *menziesii*) seed planning unit (SPU) which represents elevation bands 0-900m in geographic areas west of the British Columbia (BC) coastal mountain range and latitudinal gradients South 48°00’ - 52°00’. The CDF breeding program is the most advanced in BC and is currently in its third generation with advanced generation seed orchards producing 6.6 million seedling equivalents from 2012 to 2017. Using 14 experimental trials planted in different environments, we used monthly averages of ECs obtained from ClimateBC (Wang *et al*. 2012b), and ssGBLUP (Legarra *et al*. 2009; Misztal *et al*. 2009) under a Bayesian mixed-model framework (Pérez and de los Campos 2014) to obtain genetic parameter estimates and genomic estimated breeding values for genotyped individuals. We then used the method of Lourenco *et al*. (2015) to obtain indirect predictions of breeding values of target populations in unobserved environments. Thus, the objectives of this study were to assess the influence of including monthly average ECs for environment and G×E effects in mixed model analysis on i) genetic parameter estimates for the studied population, ii) model fit criteria, and iii) between-environment genomic prediction accuracies for intra- and inter-generational predictions.

## MATERIALS AND METHODS

### Study populations

#### Study population 1 (SP1)

The trees in this study are from low elevation (0-900m asl), third generation coastal Douglas-fir (*Pseudotsuga menziesii* (Mirb.) Franco var. *menziesii*) breeding population located in coastal British Columbia, Canada (Table 1). The parental (P0) generation consists of 78 wild plus-tree selections, which were crossed in partial disconnected diallels to produce 165 full-sib families for the second generation (F1). Full-sib families of the second generation were planted in 1975 using nursery container stock in ten environments using randomized complete block designs with four replicates per environment and four tree family row plots within the replicates (mean ≈ 15; range = 14-16; trees / full-sib family / environment). The third generation (F2) contains two series (F2-2, F2-3), with no common parentage between them (see Figure S1,Table S1, Table S2) and are the result of crossing forward selections from the F1 generation, based on tree volume. F2-2 and F2-3 were planted in 2003 and 2006 respectively, both using nursery container stock, and both planted as full-sib block tests with 5 × 5 tree plots on two environments per series (mean ≈ 19; range ≈ 17-21; trees / full-sib family / environment). A total of 31,999 tree height [cm] phenotypic measurements were available for ages 12 (F1) and 11 (F2-2 and F2-3). Phenotypes were standardized by dividing total tree height by age [years] at time of measurement to provide the mean annual increment (MAI [cm]).

**Table 1:** Experimental population summaries for pedigree-based analyses.

| Environment | N<sub>tree</sub> | N<sub>Par</sub> | N<sub>FSFam</sub> | Age | Latitude | Longitude | Elevation (mas) | Date |
| --- | --- | --- | --- | --- | --- | --- | --- | --- |
| Parents (F1) |  |  |  |  |  |  |  |  |
| Lost | 2,464 | 78 | 165 | 12 | 49° 22' 15" | 122° 14' 07" | 424 | 1975-1987 |
| Adam | 2,368 | 78 | 165 | 12 | 50° 24' 42" | 126° 09' 37" | 576 | 1975-1987 |
| Fleet | 2,595 | 78 | 165 | 12 | 48° 39' 25" | 128° 05' 05" | 561 | 1975-1987 |
| Sechelt | 2,472 | 78 | 165 | 12 | 49° 28' 05" | 123° 42' 00" | 227 | 1975-1987 |
| Eldred | 2,389 | 78 | 165 | 12 | 50° 08' 51" | 124° 11' 19" | 148 | 1975-1987 |
| Menzies | 2,509 | 78 | 165 | 12 | 50° 08' 55" | 125° 38' 16" | 333 | 1975-1987 |
| Sproat | 2,485 | 78 | 165 | 12 | 49° 17' 44" | 125° 03' 10" | 318 | 1975-1987 |
| Squamish | 2,477 | 78 | 165 | 12 | 50° 11' 51" | 123° 22' 40" | 470 | 1975-1987 |
| Tansky | 2,607 | 78 | 165 | 12 | 48° 27' 52" | 124° 01' 48" | 545 | 1975-1987 |
| White | 2,503 | 78 | 165 | 12 | 50° 06' 03" | 126° 04' 54" | 409 | 1975-1987 |
| Series 2 (F2-2) |  |  |  |  |  |  |  |  |
| Jordan | 1,551 | 74 | 79 | 11 | 48° 25' 86" | 124° 01' 80" | 160 | 2003-2014 |
| North Arm | 1,583 | 75 | 81 | 11 | 48° 50' 72" | 124° 06' 53" | 168 | 2003-2014 |
| Series 3 (F2-3) |  |  |  |  |  |  |  |  |
| Big Tree | 2,228 | 73 | 108 | 11 | 50° 15' 80" | 125° 43' 59" | 225 | 2006-2017 |
| Roach | 1,768 | 73 | 104 | 11 | 48° 45' 08" | 124° 06' 14" | 230 | 2006-2017 |

NOTE: N<sub>tree</sub>, Number of individual tree observations; N<sub>Par</sub>, Combined number of male and female parents; N<sub>FSFam</sub>, Number of full-sibling families; Age, Age of tree height measurement; Date, Year of planting to year of tree height measurement.

#### Study population 2 (SP2)

A subset of SP1 was used for the genomic analyses in this study (Table 2). The subset population was created by setting a relatedness threshold of greater than 0 as the minimum expected additive genetic kinship coefficient value (***A***, derived via pedigree) of an individual tree in the F1 or F2 generations, with at least one of the genotyped individuals in the F2 population. This resulted in 11,759 F1 phenotypes available for the genomic analyses (mean ≈ 15; range ≈ 14-16; trees / full-sib family / environment). The relatedness restriction resulted in an average additive genetic kinship coefficient of 0.033, as opposed to 0.012 in the population described previously.

**Table 2:** Training (T) and validation (V) population summaries for genomic analyses.

| Environment | Generation | N<sub>Tree</sub> | N<sub>Par</sub> | N<sub>FSFam</sub> | N<sub>Genotyped</sub> | GA1 | GA2 |
| --- | --- | --- | --- | --- | --- | --- | --- |
| Lost | F1 | 1,167 | 36 | 78 | 288 | T / V | T |
| Adam | F1 | 1,114 | 36 | 78 | 285 | T / V | T |
| Fleet | F1 | 1,228 | 36 | 78 | 295 | T / V | T |
| Sechelt | F1 | 1,164 | 36 | 78 | - | T | T |
| Eldred | F1 | 1,130 | 36 | 78 | - | T | T |
| Menzies | F1 | 1,191 | 36 | 78 | - | T | T |
| Sproat | F1 | 1,170 | 36 | 78 | - | T | T |
| Squamish | F1 | 1,169 | 36 | 78 | - | T | T |
| Tansky | F1 | 1,236 | 36 | 78 | - | T | T |
| White | F1 | 1,190 | 36 | 78 | - | T | T |
| Jordan | F2-2 | 171 | 9 | 8 | 79 | - | V |
| Bigtree | F2-3 | 821 | 26 | 41 | 230 | - | V |

NOTE: GA1, Genomic Analysis 1; GA2, Genomic Analysis 2; T, training population; V, validation population; N<sub>Tree</sub>, Number of individual tree observations; N<sub>Par</sub>, Combined number of male and female parents; N<sub>FSFam</sub>, Number of full-sibling families.

### SNP genotyping

SNP genotypes were based on those produced and used by Thistlethwaite *et al*. (2017, 2019) with the full details available in Thistlethwaite *et al*. (2017). Briefly, DNA was extracted from cambial tissue and sent to RAPiD Genomics© (Florida, US) for SNP genotyping using whole exome capture. The SNPs used here differed from those used by Thistlethwaite *et al*. (2017, 2019) due to filtering criteria resulting in a different number of SNPs and genotyped trees for this study. Filtering criteria was done using VCFtools (Danecek *et al*. 2011) and were as follows: maximum number of alleles = 2, minimum number of alleles = 2, Hardy-Weinberg Equilibrium exact test = 0.10, maximum sample missing = 0.40, maximumSNP missing = 0.40, minor allele frequency = 0.05, maximum site read depth = 50, minimum site read depth = 4. We then used the ‘impute.svd’ function from the R package ‘bcv’ (Perry 2015) to impute the remaining missing data resulting in a final count of 66,969 SNPs for the genomic analyses.

### Environmental covariates (ECs)

Thirteen monthly ECs were obtained using ClimateBC software version 5.51 (Wang *et al*. 2012b) resulting in 156 individual environmental covariates per environment (*i.e*., 12 months by 13 variables). ClimateBC generates “scale-free” climate data, thus allows the user to obtain monthly climate variables for specific test environments rather than pixel averages from grid-based climate data. Monthly ECs were averaged across the growing period for each trial, from planting year until the year of phenotypic measurement, and included primary measures of temperature, precipitation, and solar radiation (see Table S3 for a complete list of the thirteen primary and derived monthly variables).

### Pedigree-based analyses (PBLUP)

PBLUP analyses were used to estimate (co)variance components or functions of them (*i.e*., heritabilities and Type-B genetic correlations) and breeding values for the experimental population outlined in Table 1. All PBLUP analyses were completed using ASReml-R v3.0 (Butler *et al*. 2009), which uses the Average Information algorithm on Restricted Maximum Likelihood (REML) (Gilmour *et al*. 1995).

Individual-tree breeding values were obtained under the following multi-environmental mixed linear model:

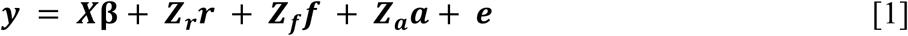

where ***y*** is the vector of phenotypic observations; **β** is the fixed effect of environment means; ***r*** is the vector of random replicate effects within of each F1 environment, with ***r***~***N***(**0, *G_r_***⊗***I***), where ***G_r_*** is the diagonal (co)variance matrix between environments with diagonal elements 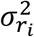. for each environment í representing the replicate within environment variances, ***I*** is the identity matrix, and ⊗ represents the Kronecker product of matrices; ***f*** is the vector of random full-sib family genetic effects, with ***f***~***N***(**0,*G_f_***⊗***I***), where ***G_f_*** is the diagonal dominance (co)variance matrix between environments with diagonal elements 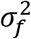 representing ¼ of dominance genetic variance (*i.e*., a common family variance for all environments was used); ***a*** is the vector of additive genetic effects (or breeding values, EBV), with ***a***~***N***(**0, G_a_**⊗***A***) where ***G_a_*** is the additive genetic (co)variance matrix between environments with diagonal elements 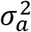 representing additive genetic variance and off-diagonal elements *σ_a_* (*i.e*., a common additive genetic variance and covariance for all environments was used) representing the same additive genetic covariance between environments, and the matrix ***A*** contains the additive genetic relationships among all trees; ***X, Z_r_, Z_f_***, and ***Z_a_*** are the respective incidence matrices assigning fixed and random effects to each observation. Finally, ***e*** is the vector of random residual effects, with ***e***~***N***(**0, *R***), where 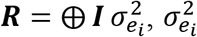 is the residual variance for each environment *i*, and ⊗ represents the ‘direct sum’ of matrices. We chose to use a common variance for genetic dominance and additive effects to allow a more parsimonious model when using a large number of trees and environments.

Single-environment narrow-sense heritability for each environment 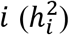 was calculated using the same model from Eq. [1], except that in ***G_a_*** different 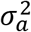 were used for each environment 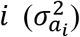 (*i.e*., CORUH structure in ASReml). Then, 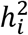 were estimated as: 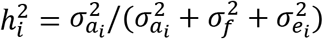.

Finally, pairwise Type-B genetic correlations between environments *i* and *i*′ (*r_gii′_*) were estimated using the model of Eq. [1] but fitting two environments at the time, and ***G_a_*** was a heterogeneous (co)variance matrix with different additive genetic variances among environments and an additive genetic covariance between pairs of environments *i* and *i*′ equal to *σ_a_ii′__*, (*i.e*., US structure in ASReml) was used for the vector random additive genetic effects in this case. For this analysis, only two environments were fitted to the model at a time. Then, *r_gii′_* was estimated as: 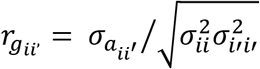.

### Genomic-based analyses (ssGBLUP)

Variance components, genetic parameters, and model fit (Deviance Information Criterion, Spiegelhalter *et al*. 2002) were estimated by fitting models to the data set in Table 2. Following Jarquín *et al*. (2014), the models fit were a series of five linear hierarchical single-step GBLUP (ssGBLUP) models. The R package ‘BGLR’ (Pérez and de los Campos 2014) was used. For variance component estimation, all data in Table 2 was fit to each model and the Monte Carlo Markov Chain (MCMC) was generated with 200,000 iterations thinned at a rate of 5, with the first 25,000 discarded as burn-in. Bayesian ridge regression (BRR; Gaussian prior) model was used for all model effects. The ‘BGLR’ package default starting parameters were used. Multiple chains were generated and the MCMC chain posterior means and trace plots of the posterior distributions were compared to confirm convergence. Cholesky decomposition of all variance-covariance matrices was used to speed up computations.

#### Model 1 (M1)

The first ssGBLUP model included an overall mean (μ) as a fixed effect, random environment effects (***S_i_***), random replicate (***R_k_***), and random additive genetic effects (***a_j_***). Let ***y_ijk_*** and ***e_ijk_*** be the phenotype and residual effects, respectively, of individual *j* for environment *i*, scored in replicate *k*. Then the mixed model to analyze the data is:

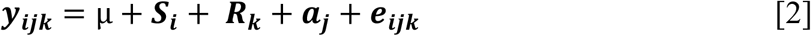

where 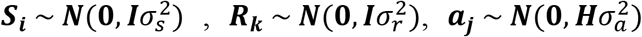 and 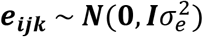 where the respective variances 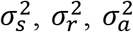 and 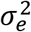 are common for all the environments. The combined additive genetic relationship matrix, ***H***, was calculated as follows (Legarra *et al*. 2009; Christensen and Lund 2010):

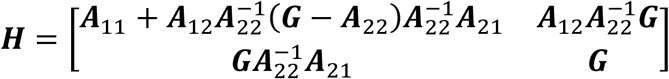

where ***A***_11_ is the portion of the average numerator relationship matrix (***A***) including the non-genotyped individuals, ***A***_22_ is the portion of genotyped individuals, and ***A***_12_ and ***A***_21_ are the portions containing the expected additive genetic relationships between genotyped and non-genotyped individuals. The genomic additive relationship matrix for genotyped individuals, ***G***, was calculated after VanRaden (2008):

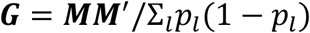

where ***M*** is the centered matrix of SNP covariates, and *p_l_* is the current (or observed) allele frequency of the genotyped trees for marker *l*. Further, ***G*** was scaled for compatibility with ***A*** such that the mean diagonal and off-diagonal elements were equal (Christensen *et al*. 2012):

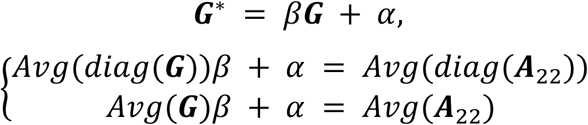

To avoid potential problems with the inversion of ***G***\* it was weighted as (Aguilar *et al*. 2010):

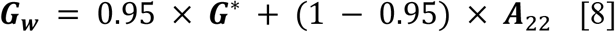

#### Model 2 (M2)

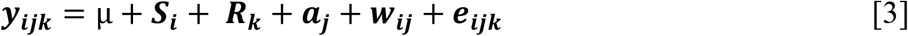

M2 extends model M1 (Eq. [2]) to include the vector of random effects for environmental covariates (***w***) which is distributed as 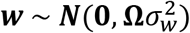. Where **Ω** is the matrix of similarity among environments calculated as **Ω** ∝ ***WW***′/*q*, with ***W*** being a centred and standardized matrix of ECs and *q* the number of ECs as described by (Jarquín *et al*. 2014) in detail, and 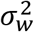 is the variance of the vector of ECs.

#### Model 3 (M3)

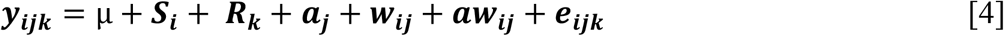

M3 incorporates the vector of random effects for the interaction between additive genetic (***a***) and 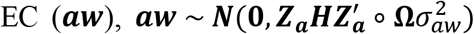, where 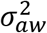 is the variance of the interaction between additive genetic effects and ECs, and ∘ is the Hadamard (Schur) product of matrices.

#### Model 4 (M4)

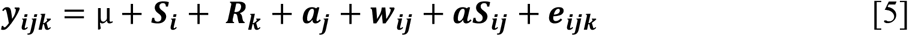

M4 includes a vector of random effects for the interaction between additive genetic and main environment terms (***aS***), 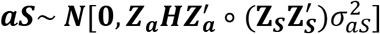, where ***Z_s_*** is the incidence matrix for the effects of environments, and 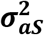 is the variance of the interaction between additive genetic and main environmental effects.

#### Model 5 (M5)

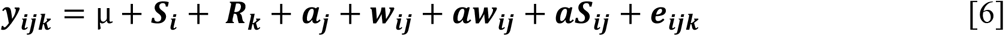

Finally, M5 includes both interaction effects from M3 (i.e., ***aw***) (Eq. [4]) and M4 (i.e., ***aS***) (Eq. [5]).

### Prediction accuracy and cross-validation

Two scenarios were considered for estimating the prediction accuracy of the validation populations (VP, Table 2). In the first scenario (GA1), a leave one environment out approach for environments with genotyped individuals (Lost, Adam, and Fleet) was used to form the training population (TP). In the second scenario (GA2), the TP consisted of all individuals from the F1 generation. In either case, the non-genotyped individuals of the TP were randomly divided into five folds, that is, the phenotypes of genotyped individuals in the TP always participated in the estimation of SNP marker effects. The random folding process was replicated three times for each model (M1-M5). For each fold, genomic estimated breeding values (GEBV) of the genotyped VP were correlated to their EBV estimated in the PBLUP analysis (Eq. [1]). Prediction accuracy (*r*_(*GEBV,EBV*)_) was then calculated as the mean Pearson product-moment correlation of the three replicates. The procedure for obtaining GEBV for the VP follow closely the methods of Lourenco *et al*. (2015) and are outlined below.

### Estimation of GEBV for the validation population

1. Use ssGBLUP models M1-M5 to obtain the vector of GEBV (***a***) for the genotyped portion of the TP.
2. Calculate the direct genomic value for each genotyped tree *j*(*DGV_j_*) in the TP (Aguilar et al. 2010):

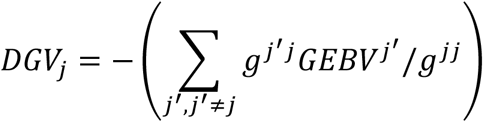

where *g^j’j^* (and *g^jj^*) is an off-diagonal (and diagonal) element of the inverse of ***G***(***G^−1^***) corresponding to relationships between tree *j′* and *j* and *GEBV^j′^* is the predicted additive genetic solutions from the ssGBLUP models M1-M5 for TP and individual *j′*.
3. Estimate SNP marker effects from the TP:

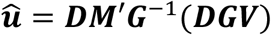

where ***û*** is the vector of estimated SNP effects, ***D*** is a weight diagonal matrix for SNP (here an identity matrix), ***M*** as defined before, and DGV is the vector of DGV for the TP.
4. Calculate DGV for the trees in the VP:

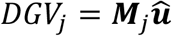

where *DGV_j_* and ***M***_*j*_ are the direct genomic values and the matrix of centered genotypes for tree *j* in the VP and not included in the ssGBLUP models M1-M5, respectively.
5. Calculate GEBV for the VP:

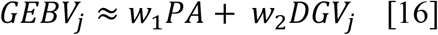

where PA is the parental average of individual *j* in the VP and *w*_1_ and *w*_2_ are weights associated with the covariances of *DGV_j_* and PA:

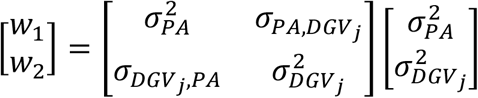 Lourenco *et al*. (2015) note that this approximation excludes the pedigree prediction component from the selection index approach proposed by VanRaden *et al*. (2009). Parental breeding values for the F1 generation were calculated as the average breeding value of all offspring without masked phenotypes. Parental breeding values for the F2 generation were directly estimated in the prediction models.

### Data availability

All statistical analyses were carried out in the R environment (R Core Team 2017). Phenotypic and genotypic data are available from the Dryad Digital Repository https://doi.org/10.5061/dryad.8n2d374.

## RESULTS

### Pedigree-based analyses (PBLUP)

Individual tree single environment narrow-sense heritability estimates for the PBLUP analysis were low to moderate for the F1 generation trials 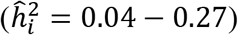 while estimates for the F2 generation were higher 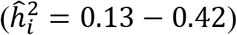 (Table 3). Standard errors of heritability estimates were low indicating significance of the estimates. Type-B genetic correlation estimates between F1 trials were positive and generally high (*r̂_gii′_* = 0.51 − 0.98), representing agreement in allelic effects contributing to MAI among the tested environments. This result is expected since the trials are from a single breeding zone. Among the three genotyped F1 trials (Lost, Adam, Fleet) the Type-B genetic correlations were moderate to high (*r̂_gii′_* = 0.71 − 0.88), and between environments of F2-2 (*r̂_gii_* = 0.62) and F2-3 (*r̂_gii′_* = 0.57). Type-B genetic correlations between F1 and F2 trials were in most cases not significant based on standard errors or were non-estimable due to low genetic connectivity between environments (see Figure S1, Table S1, Table S2).

**Table 3:** Pedigree based single environment narrow-sense heritabilities (diagonal) and Type-B genetic correlations (lower triangle) for MAI, standard errors in parentheses.

|  | Lost | Adam | Fleet | Sechelt | White | Menzies | Sproat | Tansky | Eldred | Squamish | Bigtree | Roach | Jordan | Northarm |
| --- | --- | --- | --- | --- | --- | --- | --- | --- | --- | --- | --- | --- | --- | --- |
| Lost | **0.13 (0.002)** |  |  |  |  |  |  |  |  |  |  |  |  |  |
| Adam | 0.71 (0.128) | **0.18 (0.002)** |  |  |  |  |  |  |  |  |  |  |  |  |
| Fleet | 0.83 (0.131) | 0.88 (0.089) | **0.22 (0.004)** |  |  |  |  |  |  |  |  |  |  |  |
| Sechelt | b | 0.77 (0.097) | 0.98 (0.071) | **0.22 (0.003)** |  |  |  |  |  |  |  |  |  |  |
| White | 0.73 (0.138) | 0.85 (0.100) | 0.93 (0.078) | 0.93 (0.078) | **0.04 (0.014)** |  |  |  |  |  |  |  |  |  |
| Menzies | 0.83 (0.137) | 0.90 (0.095) | 0.92 (0.117) | 0.91 (0.099) | 0.97 (0.097) | **0.17 (0.003)** |  |  |  |  |  |  |  |  |
| Sproat | 0.55 (0.163) | 0.51 (0.142) | 0.78 (0.116) | 0.80 (0.104) | 0.78 (0.115) | 0.81 (0.128) | **0.21 (0.003)** |  |  |  |  |  |  |  |
| Tansky | 0.73 (0.138) | 0.63 (0.121) | 0.89 (0.083) | 0.84 (0.090) | 0.67 (0.125) | 0.72 (0.132) | 0.60 (0.136) | **0.27 (0.005)** |  |  |  |  |  |  |
| Eldred | 0.77 (0.159) | 0.61 (0.148) | 0.77 (0.144) | 0.98 (0.100) | 0.91 (0.108) | 0.89 (0.139) | 0.86 (0.139) | 0.72 (0.148) | **0.14 (0.002)** |  |  |  |  |  |
| Squamish | 0.83 (0.103) | 0.85 (0.068) | 0.87 (0.076) | 0.84 (0.080) | 0.80 (0.090) | 0.93 (0.070) | 0.62 (0.120) | 0.64 (0.120) | 0.70 (0.135) | **0.25 (0.002)** |  |  |  |  |
| Bigtree | -0.23 (0.604) | -0.01 (0.474) | 0.53 (0.394) | -0.14 (0.472) | 0.09 (0.529) | 0.13 (0.597) | -0.53 (0.455) | -0.04 (0.554) | -0.06 (0.575) | -0.05 (0.565) | **0.13 (0.000)** |  |  |  |
| Roach | 0.48 (0.468) | 0.46 (0.368) | 0.32 (0.418) | 0.31 (0.407) | 0.20 (0.471) | 0.51 (0.452) | -0.03 (0.506) | 0.24 (0.480) | 0.19 (0.500) | 0.70 (0.338) | 0.57 (0.133) | **0.33 (0.000)** |  |  |
| Jordan | 0.73 (0.488) | 0.64 (0.430) | 0.20 (0.504) | 0.72 (0.400) | 0.80 (0.540) | b | 0.98 (0.369) | 0.41 (0.573) | b | 0.35 (0.455) | b | b | **0.42 (0.006)** |  |
| Northarm | b | b | b | 0.89 (0.430) | b | b | b | b | b | 0.75 (0.435) | b | b | 0.62 (0.194) | **0.12 (0.003)** |

NOTE: b, Type-B genetic correlation and their approximate standard errors were not estimated due to convergence problems.

### Genomic-based analyses (ssGBLUP)

In M1 46% and 5% of the phenotypic variance was explained by the between environment 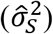 and within environment 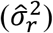 effects, respectively (Table 4). With the addition of ECs 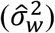 in M2 those values dropped to 34% and 3% respectively, with the ECs accounting for 22% of the phenotypic variance. The shift of explained variance from the between environment effect to ECs can be understood as a decomposition of environmental variance, showing that ECs in M2 are able to capture additional environmental variation not captured by M1. This shift was accompanied by 9% decrease in error variance 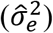 across models M1-M5. However, a minor increase in the DIC model fit statistic of M2 did not justify the addition of ECs to M1.

**Table 4:** Model Deviance Information Criterion (DIC), posterior means and their 95% highest posterior density (HPD) intervals in brackets for the additive variance 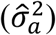, replicate within environment variance 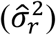, environmental variance 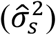, environment covariates variance 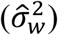, the interaction between additive effects and environment covariates 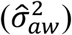, the variance of the interaction between additive and environmental effects 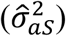, residual variance 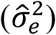, heritability (*ĥ*^2^). See text for models’ abbreviations.

| Model | DIC | $\hat{\sigma}_a^2$ | $\hat{\sigma}_r^2$ | $\hat{\sigma}_s^2$ | $\hat{\sigma}_w^2$ | $\hat{\sigma}_{aw}^2$ | $\hat{\sigma}_{as}^2$ | $\hat{\sigma}_e^2$ | $\hat{h}^2$ |
| --- | --- | --- | --- | --- | --- | --- | --- | --- | --- |
| M1 | 87846.80 | 20.02<br>[13.00 27.59] | 11.11<br>[6.35 16.63] | 102.53<br>[38.18 193.00] | -<br>- | -<br>- | -<br>- | 91.35<br>[86.24 96.17] | 0.18<br>[0.12 0.24] |
| M2 | 87850.41 | 19.11<br>[12.58 26.08] | 9.20<br>[5.27 14.01] | 90.52<br>[24.32 186.18] | 58.04<br>[10.15 181.48] | -<br>- | -<br>- | 91.55<br>[86.67 96.23] | 0.17<br>[0.12 0.23] |
| M3 | 87176.25 | 18.51<br>[11.89 25.15] | 8.05<br>[4.65 12.22] | 86.16<br>[26.04 168.63] | 33.86<br>[8.22 70.70] | 19.99<br>[14.88 25.31] | -<br>- | 76.61<br>[71.10 82.19] | 0.16<br>[0.11 0.21] |
| M4 | 87380.72 | 17.41<br>[11.27 23.88] | 8.03<br>[4.57 12.32] | 85.50<br>[21.92 172.05] | 38.95<br>[8.84 99.37] | -<br>- | 12.82<br>[9.67 15.94] | 81.12<br>[76.17 86.00] | 0.16<br>[0.10 0.21] |
| M5 | 87159.53 | 16.53<br>[10.87 22.92] | 7.20<br>[4.07 10.97] | 90.08<br>[25.45 181.30] | 30.08<br>[5.37 65.72] | 12.19<br>[6.96 17.48] | 8.52<br>[5.64 11.49] | 76.43<br>[71.27 81.92] | 0.15<br>[0.10 0.20] |

Evaluating the addition of G×E interactions into the model structure (M3-M4) showed that the use of ECs 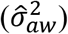 in M3 explained 4% more phenotypic variation than the use of the environment interaction effect 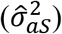 in M4. Large decreases in the DIC model fit statistic were observed with the addition of either ***aS_ij_*** or ***aw_ij_*** interaction effects in the model, justifying their use; however, when considering them individually the ***aw_ij_*** effect offered a better DIC model fit statistic. The joint use of both ***aS_ij_*** and ***aw_ij_*** interaction effects in the model to explain G×E accounted for 9% of the total phenotypic variation, indicating a small but important source of genetic variation in the breeding population. The DIC estimate for the most complex model (M5) was lowest, justifying the specification and use of all model parameters for the prediction of breeding values in this population.

Narrow-sense heritability estimates for models M1-M5 were in agreement with the mean single environment narrow-sense heritability estimates from the prior PBLUP analysis. A decrease in additive genetic variance estimates and consequently narrow-sense heritability, was observed with increasing model complexity due primarily to the addition of G×E effects into the models, and a decrease in percent variance explained by the additive genetic effect 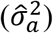 9% M1 vs 7% M2-M5. The decrease in heritability highlights the confounding nature between additive genetic effect estimates and G×E interaction effects.

### Genomic prediction accuracy

#### Intra-generation (GA1)

Models M1-M5 prediction accuracies for EBV of genotyped individuals across environments and within the F1 generation varied (*r*_(*GEBV,EBV*)_ = 0.447 − 0.640) (Table 5, Figure 1). Individual environment mean prediction accuracies for models M1-M5 were not equal among the three tested environments *r*_(*GEBV,EBV*)_ = 0.619 (Lost), *r*_(*GEBV,EBV*)_ = 0.513 (Fleet), and *r*_(*GEBV,EBV*)_ = 0.452 (Adam). The addition of ECs facilitated by model M2 gave minor increases in prediction accuracy, up to 2% over M1 (Fleet). Gains in prediction accuracy of up to 6% (M3, Lost; M5, Fleet) were observed by accounting for G×E effects in the population. Treatment of G×E effects using ECs (M3) performed better than using main environment effects (M4) when comparing to the base model scenario (M2). Standard errors of predictions were minor, ranging from 0.0000-0.0027, indicating low variation between cross-validation replicates.

**Figure 1:**
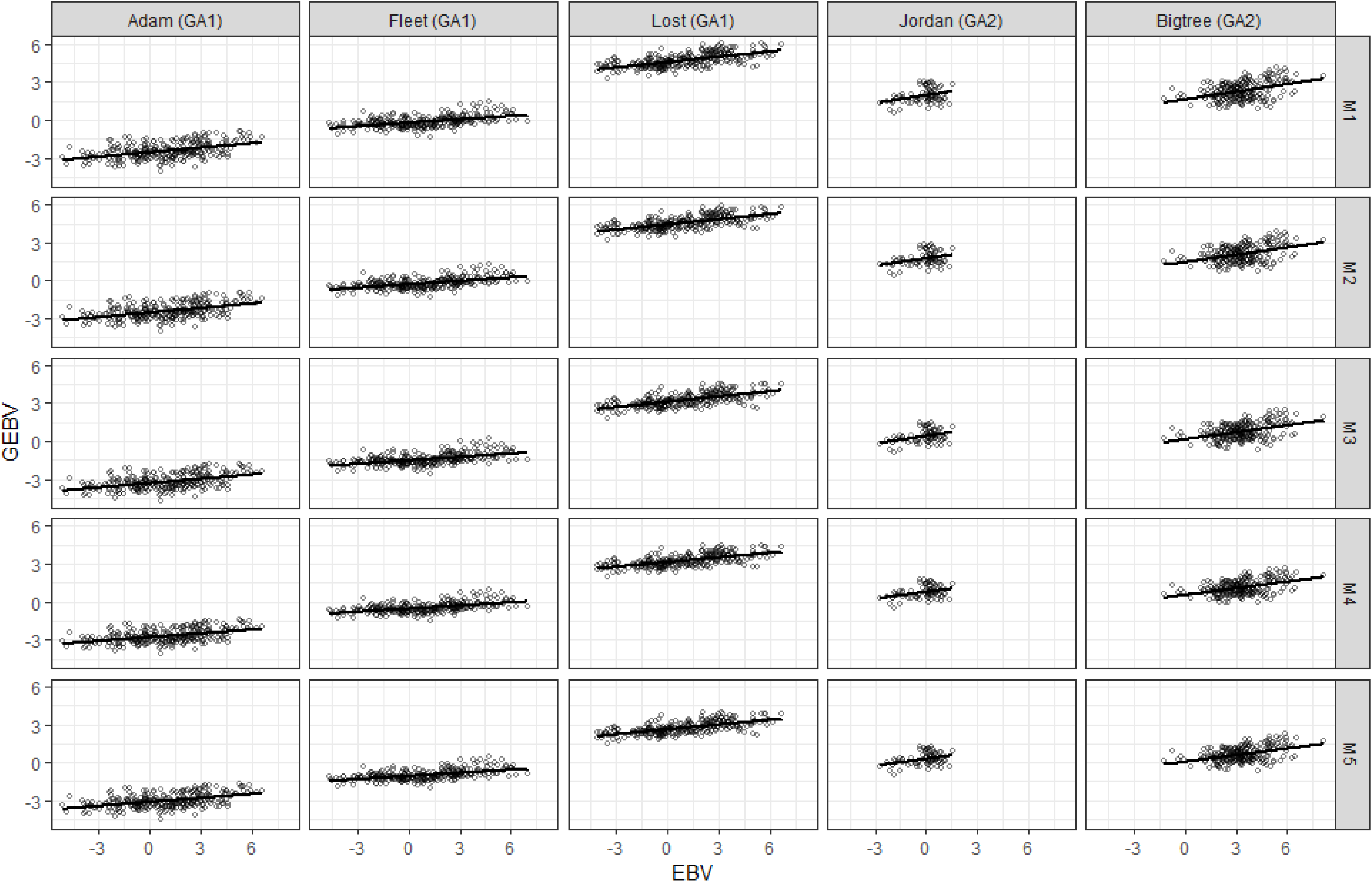
Scatterplots of intra-generation (GA1) and inter-generation (GA2) predictions of EBV.

**Table 5:** Intra-generation (GA1) cross-validation prediction accuracies (*r*_(GEBV,EBV)_) with standard errors (*SE*). See text for models’ abbreviations.

|  | Lost |  | Adam |  | Fleet |  |
| --- | --- | --- | --- | --- | --- | --- |
| | $r_{(GEBV,EBV)}$ | $SE$ | $r_{(GEBV,EBV)}$ | $SE$ | $r_{(GEBV,EBV)}$ | $SE$ |
| M1 | 0.601 | 0.0000 | 0.447 | 0.0004 | 0.498 | 0.0006 |
| M2 | 0.613 | 0.0011 | 0.453 | 0.0011 | 0.513 | 0.0027 |
| M3 | 0.640 | 0.0007 | 0.461 | 0.0006 | 0.519 | 0.0010 |
| M4 | 0.615 | 0.0011 | 0.448 | 0.0010 | 0.515 | 0.0010 |
| M5 | 0.629 | 0.0009 | 0.452 | 0.0010 | 0.526 | 0.0023 |

#### Inter-generation (GA2)

Testing the models to predict EBV across generations and across environments yielded an expectedly lower minimum prediction accuracy than observed for GA1 (*r*_(*GEBV, EBV*)_ = 0.317 − 0.538) (Table 6, Figure 1). In all models, prediction accuracies were greater for the Bigtree environment than they were for Jordan. Comparison of models M1 and M2 show that the addition of ECs in M2 did not produce an increase in prediction accuracy for Bigtree and a 1% decrease was observed for Jordan. The addition of individual G×E terms in models M3 and M4 both gave increases in prediction accuracy over the base model M1 for both Jordan and Bigtree. However, the use of ECs to explain G×E variation (M3) gave 13% (Jordan) and 6% (Bigtree) increases in prediction accuracies over M1 versus 8% (Jordan) and 4% (Bigtree) given by M4. No improvement in prediction accuracy occurred for Jordan with the addition of both G×E terms in model M5 and estimates were, in fact, equal to those given by M3. For the Bigtree environment, a small 2% gain in prediction accuracy was observed for model M5 over model M4 versus 1% for model M3.

**Table 6:** Inter-generation (GA2) cross-validation prediction accuracies (*r*_(*GEBV,EBV*)_) and standard errors (*SE*). See text for models’ abbreviations.

|  | Jordan |  | Bigtree |  |
| --- | --- | --- | --- | --- |
| | $r_{(GEBV,EBV)}$ | $SE$ | $r_{(GEBV,EBV)}$ | $SE$ |
| M1 | 0.320 | 0.0012 | 0.506 | 0.0006 |
| M2 | 0.317 | 0.0011 | 0.506 | 0.0008 |
| M3 | 0.360 | 0.0015 | 0.535 | 0.0009 |
| M4 | 0.346 | 0.0029 | 0.526 | 0.0006 |
| M5 | 0.360 | 0.0016 | 0.538 | 0.0004 |

## DISCUSSION

The capability of GS and its application to increase genetic gain in conifer forest tree breeding programs is at this point recognized as being a future certainty (Grattapaglia *et al*. 2018). Genetic gain is ultimately a product of trait heritability, accuracy, and intensity of selection, and the time interval to complete a round of selection, testing, and breeding. Of these factors, the accuracy of selection and time effort are most effectively addressed by GS methods. Previously, with forest trees, the ssGBLUP method was demonstrated by Ratcliffe *et al*. (2017), Cappa *et al*. (2018) and Klápště *et al*. (2018) to improve the accuracy of genetic parameter estimates and intra-generation breeding values in forest trees. Here, the ssGBLUP methodology permitted the use of rogued genetic trial data by collectively leveraging their phenotypic, pedigree and genomic information to participate in marker effect estimation and subsequent intra- and inter-generation breeding value predictions. Additionally, the merit of using indirect genomic predictions with the ssGBLUP method was supported by this study in reducing the computational time required to obtain predictions for unobserved samples.

We further demonstrated the capabilities of ssGBLUP in boosting (or increasing) the training population size without the need of extra genotyping, an important aspect of genomic prediction accuracy highlighted by Hayes et al. (2009). As a result, the training population (*n* = 11,759) used here, so far represents the largest for a genomic prediction study in forest trees. A study involving Norway spruce highlighted that a TP to VP ratio of up to 3:1 produced increased prediction accuracies for growth and wood quality traits (Chen *et al*. 2018). However, it should be noted that Tan et al. (2017) and Lenz et al. (2017) observed benefit in prediction accuracy above the stated 3:1 ratio. Training population size is critical when the G×E effect can be as substantial as commonly observed in forest trees. Thus, it is a boon when historical trials which represent testing of genotypes across many environments can be recovered and included in the genomic prediction analysis. Additionally, larger training populations will better capture trait architecture and the totality of genetic diversity of the breeding population, a substantial benefit for making inferences. This variation may include rare marker variants, which are effective for the prediction of oligogenic traits where they may account for larger proportions of trait heritability (e.g., Resende *et al*. 2017). In our analysis, the studied trait (MAI) is considered to have low heritability and complex genetic architecture (*i.e*., polygenic). Indeed, the heritability estimates (Table 3, Table 4) and marker effect plot (Figure S2) agree, justifying the use of the BRR model and Gaussian prior choice for prediction of the GEBV.

We investigated the effect of modeling G×E interactions on GS prediction accuracies in Douglas-fir, an outcrossing conifer tree species. Thereby, we demonstrated increased inter- and intra-generation prediction accuracy of EBV through consideration of G×E effects defined not only by main environment effects but also by ECs in the predictive models. The inclusion of EC effects in the model by themselves (*i.e*., M2) presented a fractioning of the phenotypic variance into effects explained by the main environment effect (34%) and that of the ECs (22%). This was accompanied by a 7% decrease in the proportion of total variance explained by the residual effect, thus producing more realistic gain estimates. However, in most cases M2 offered little to no increase in prediction accuracy over M1, which agrees with the results of previous studies which tested the same suite of models used here for various traits in plants such as rice, wheat, or cotton (Jarquín *et al*. 2014; Pérez-Rodríguez *et al*. 2015; Morais Júnior *et al*. 2018). Jointly, the specification of EC effects, ***w_ij_*** and ***aw_ij_***, in M3 resulted in a decrease in estimated residual variance over M1 (16%) and the interaction effect explained 9% of the phenotypic variation versus 5% of ***aS_ij_***, the interaction specified using the main environment effect (M4). Additionally, models M3 and M5 showed the most consistent increases in prediction accuracy over the base model M1 which also corroborates the aforementioned crop plant-based studies.

In our analysis, maximum intra-generation prediction accuracies were achieved using either M3 or M5 which always included ***w_ij_*** and ***aw_ij_*** and ranged from *r* = 0.461 − 0.640 depending on the target environment, which are similar to previous reports of intra-generation GS prediction accuracies based on multi-environment trials using conifer trees. Though many previous studies did not explicitly model or test the inclusion of G×E interaction effects in the training population or GS prediction model. For example, and perhaps most informative, was a four environment GS analysis of loblolly pine (*Pinus taeda*) by Resende *et al*. (2012) which showed an average decrease of 23% in prediction accuracies for tree height (age 6) when the prediction was done in unobserved environments within the same breeding zone (*r* = 0.41 − 0.74). However, the same study noted an average decrease of 41% when the prediction was extended to between breeding zones (*r* = 0.23 − 0.55). Thus, Resende *et al*. (2012) stressed the importance of regionalizing tree breeding efforts, breeding zone delineation, and the lack of transferability of genomic prediction models across breeding zones. A later study by Beaulieu *et al*. (2014) found large decreases (33-41%) in prediction accuracies for Quebec white spruce (*Picea glauca*) tree height (age 17) when predicting EBV in unobserved environments (*r* = 0.22 − 0.34). Lenz (2017) however, reported only 2% and 9% reductions in prediction accuracy for 25 year tree height for black spruce (*Picea mariana*) when their prediction model did not include the target environment (*r* = 0.52 − 0.56). The results observed by Lenz *et al*. (2017) reflect the low G×E effects of their black spruce population.

Two previous studies did however explicitly model GxE in the training population. The first by Gamal El-Dien *et al*. (2015) showed an average decrease of 29% in genomic prediction accuracy of tree height (age 38) for interior spruce (*Picea glauca* × *engelmannii*) when the target environment was not explicitly included in the training population (*r* = 0.37 − 0.53). However, the mentioned study did not compare the inclusion of specific model terms for G×E to investigate their impact on prediction accuracy as was done in the present study. The second by Thistlethwaite *et al*. (2017) was based on the same F1 population of trees studied here, albeit with differences (see further discussion). Thistlethwaite *et al*. (2017) observed prediction accuracies of *r* = 0.88 − 0.93 for age 12 tree height. This result contrasts with previous studies, since it is higher than the average prediction accuracy when the training population contained observations from all three environments (*r* = 0.88).

Empirical studies examining inter-generation GS prediction accuracies in forest tree species are very limited. Isik *et al*. (2016) were the first to include multiple generations in a genomic selection analysis of a forest tree. The prior study was continued by Bartholomé *et al*. (2016) who observed moderately high prediction accuracies (*r* = 0.70) for age 12 tree height when using a training population of grandparents and parents to predict offspring. Using samples from the same breeding population of this study, Thistlethwaite *et al*. (2019) observed inter-generation prediction accuracy of *r* = 0.42 for the Jordan environment. However, the results presented by Thistlethwaite *et al*. (2019) are not directly comparable to those presented here due to many key methodological differences: i) genomic prediction methods, ii) training and validation population structure and composition, iii) marker subset and imputation method, iv) estimation method of EBV, and v) modeling of G×E factors. The inter-generation prediction accuracies observed in this study were nevertheless lower than those of the study for maritime pine juvenile tree height (*r* = 0.70) (Bartholomé *et al*. 2016). Maximum inter-generation prediction accuracies for MAI here were *r* = 0.360 (Jordan) and *r*_(*GEBV,EBV*)_ = 0.538 (Bigtree) given by model M6 in both cases.

The inclusion of a main G×E interaction term is standard in individual-tree mixed models, yet the inclusion of ECs to explain environmental heterogeneity and genetic variation is not yet widely used in forestry. With the availability of ‘scale-free’ climate data for specific locations from programs such as ClimateBC (Wang *et al*. 2012b) and the NASA POWER Project (Stackhouse *et al*. 2017), breeders should opt for their use to help describe growing conditions and improve EBV predictions. Type-B genetic correlations in this study were high between the predicted F1 environments (*ȓ_gii′_* = 0.71 − 0.88), which might account for the moderately high intra-generation prediction accuracies observed here. However, Coastal Douglas-fir is noted to have strong G×E interaction for growth traits (Campbell 1992; Cappa *et al*. 2016), which may also help to explain the observed gain in prediction accuracy in this study by accounting for G×E effects. Thus, it appears to be critical for species with moderate to strong G×E interactions to either i) explicitly include the target environment in the GS prediction model, or ii) incorporate G×E effects in the genomic prediction model by including observations from as many environments as possible from across the breeding zone, an opportunity made possible with ssGBLUP.

To our knowledge, this is the first use of ECs to capture environmental heterogeneity and G×E effects on phenotypic variation in outcrossing trees for genomic prediction of breeding values. The prior mentioned studies concerning forest trees and multi-environment trials have not considered methods to improve predictions in unobserved environments as was done in this study. Here, we advocate the inclusion of ***w_ij_***, as well as both ***aS_ij_*** and ***aw_ij_*** interaction effects in the model as the gain in prediction accuracy was generally consistent across all tested environments and generations.

From a practical standpoint, the base (M2) and fully specified (M5) models were mostly agreeable in candidate rankings of the predicted individuals from unobserved environment (Figure S3). It is clear from the low discordance in rankings of highest and lowest ranked individuals that the same selections would be made irrespective of the model used. However, forest tree breeding programs by necessity require minimum effective population sizes (*N_e_*) which requires the selection of some sub-optimal individuals to maintain adequate genetic diversity in the breeding population. For these individuals (*i.e*., mid-ranked candidates) there is much less concordance in rankings between the two models. During this phase the selection of these individuals becomes important to balance genetic gain with and genetic diversity in the breeding population. This result is in agreement with Stejskal et al. (2018) who noted the capacity of GS to accurately capture Mendelian sampling (*i.e*., within family genetic variance) and cryptic relatedness in the breeding population, making it a preferred approach over traditional pedigree-based selection in forestry.

In addition to the combination of high-density genomic and environmental/climate data, it is realistic to assume that prediction of phenotypes can be further improved through the simultaneous inclusion of multiple phenotypic and environmental traits obtained via high-throughput platforms such as those produced by remote sensing (Araus *et al*. 2018; Dungey *et al*. 2018). Information sharing between genetic lines, correlated traits and correlated environments via multiple-trait models with the specification of G×E interaction effects will lead to improved phenotypic predictions. Finally, as the anticipated increasing role of GS in tree breeding, it is essential to highlight the importance of prediction accuracy improvement through the utilization of all possible available information such as was demonstrated here with the inclusion of non-genotyped individuals and their testing environments, as well as environmental covariates to better model G×E effects.

## ACKNOWLEDGEMENTS

The authors would like to thank Odilon Peixoto de Morais Júnior for their help with the statistical analysis.

This work was supported by Genome British Columbia (User Partnership Program (UPP-001) to YEK and MS, NSERC Discovery Grant to YEK, TimberWest Forest Corp., and Western Forest Products Inc. We declare that the funding agencies did not participate in the design of the study and collection, analysis, and interpretation of data and in writing the manuscript.

## AUTHOR CONTRIBUTIONS

BR, YEK, and MS conceived the study. BR, EPC, and FT performed the data analysis. MS, IP, FT for phenotypic data collection and DNA extraction. BR drafted the manuscript. CC, EPC, OG, IP, TW, JK, MS, YEK, and FT provided instrumental comments, feedback, and technical support.

## SUPPLEMENTS

**Figure S1:**
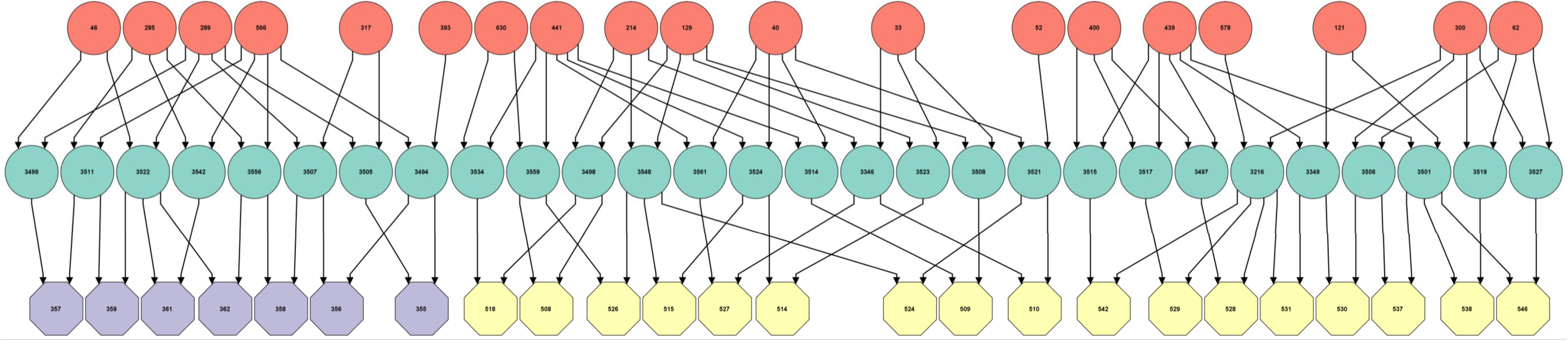
Three generation coastal Douglas-fir contemporary pedigree containing the inter-generation validation population of study. Circles represent single individuals; octagons represent full-sibling families. Red colours are the unrelated (assumed) base population of wild plus tree selections (P0), green colours are the forward selection progenitors (F1) of the Jordan (purple, F2-2) and Bigtree (yellow, F2-3) validation populations (F2) respectively. The pedigree was produced using Helium software (Shaw *et al*., 2014).

**Figure S2:**
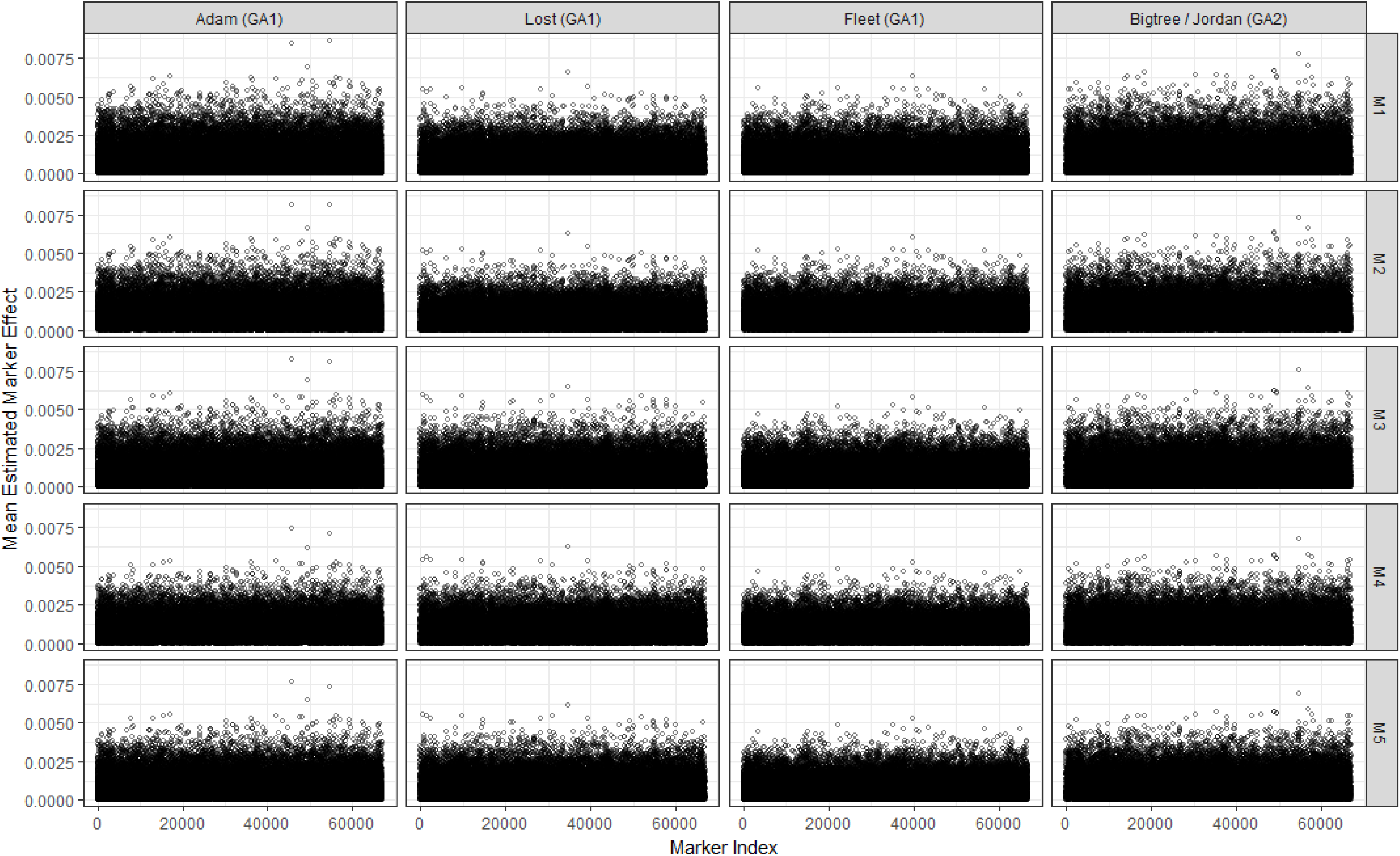
Scatterplot of mean absolute marker effects from the cross-validation analysis.

**Figure S3:**
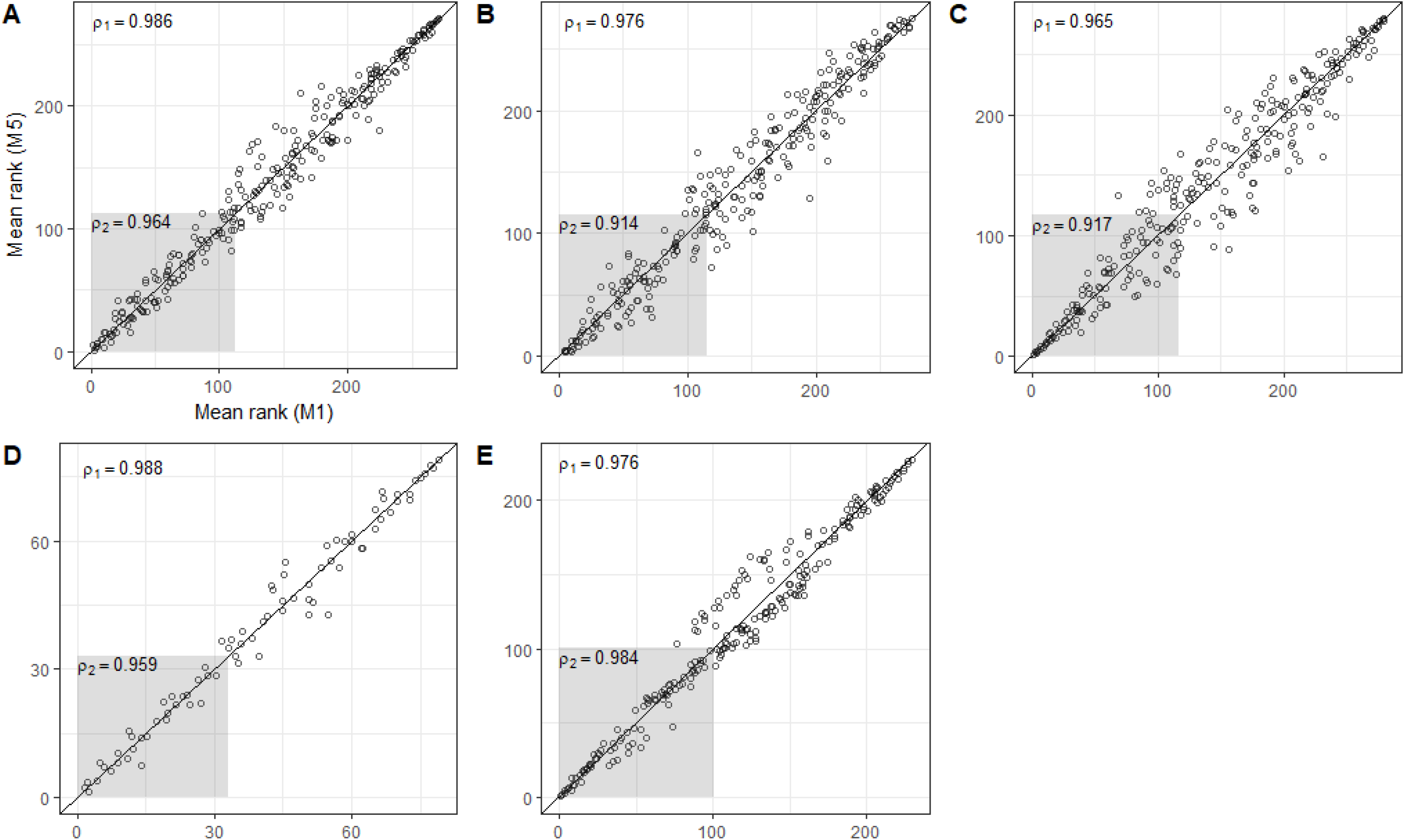
Scatterplots of mean rank of predicted trees with Spearman rank correlation coefficients from the cross-validation analyses for all predictions (*ρ*_1_) and top 40% of predictions (*ρ*_2_) within environments Adam (A), Lost (B), Fleet (C), Jordan (D), and Bigtree (E) comparing the base model (M1, x-axis) and fully specified model (M5, y-axis). Deviation from the diagonal line implies a rank change.

**Table S1:**
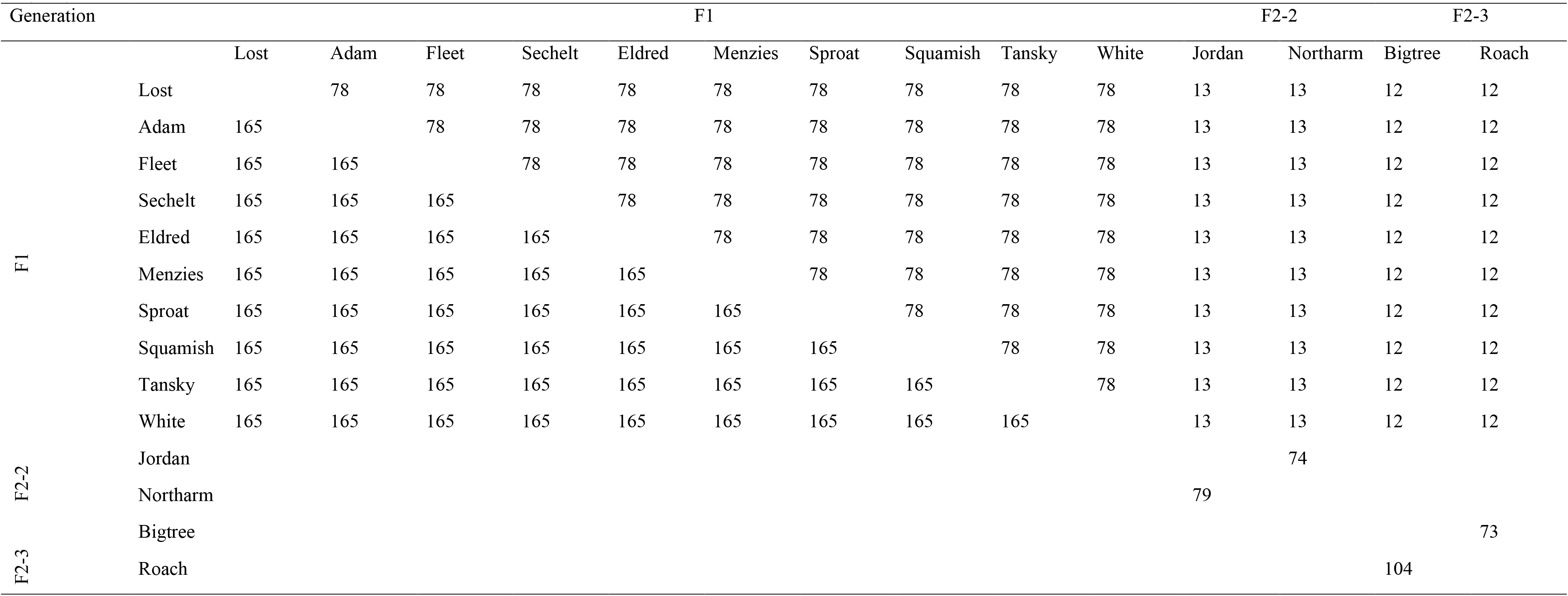
Number of parent-parent (F1-F1 / F2-F2) or parent-grandparent (F1-F2) (above diagonal) and families (below diagonal) in common among the environments for study population 1 (SP1).

**Table S2:**
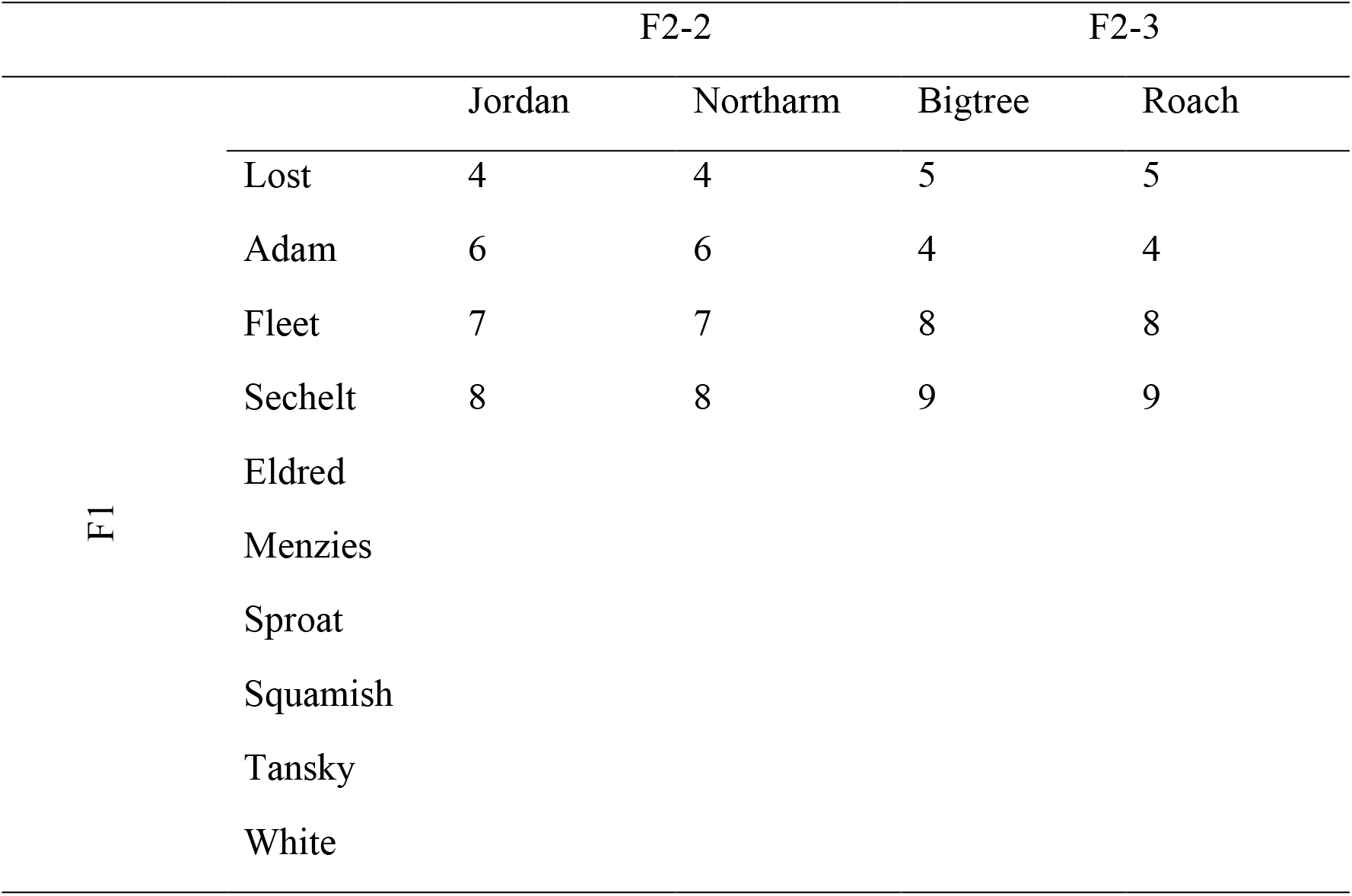
Number of F2 parents within the F1 environments.

**Table S3:** List of monthly (from January/01 to December/12) climatic variables obtained from ClimateBC v5.51 (Wang *et al*., 2012a) used in the genomic analyses.

| Primary monthly variables |  |
| --- | --- |
| Tave01 – Tave12 | mean temperatures (°C) |
| TMX01 – TMX12 | maximum mean temperatures (°C) |
| TMN01 – TMN12 | minimum mean temperatures (°C) |
| PPT01 – PPT12 | precipitation (mm) |
| RAD01 – RAD12 | solar radiation (MJ m<sup>-2</sup> d<sup>-1</sup>) |
| Derived monthly variables |  |
| DD_0_01 – DD_0_12 | degree-days below 0°C |
| DD5_01 – DD5_12 | degree-days above 5°C |
| DD_18_01 – DD_18_12 | degree-days below 18°C |
| DD18_01 – DD18_12 | degree-days above 18°C |
| NFFD01 – NFFD12 | number of frost-free days |
| PAS01 – PAS12 | precipitation as snow (mm) |
| Eref01 – Eref12 | Hargreaves reference evaporation (mm) |
| CMD01 – CMD12 | Hargreaves climatic moisture deficit (mm) |

